# Uneven protection of persistent giant kelp forest in the Northeast Pacific Ocean

**DOI:** 10.1101/2020.12.15.422778

**Authors:** Nur Arafeh-Dalmau, Kyle C. Cavanaugh, Hugh P. Possingham, Adrian Munguia-Vega, Gabriela Montaño-Moctezuma, Tom W. Bell, Kate Cavanaugh, Fiorenza Micheli

## Abstract

In most regions, the distribution of marine forests and the efficacy of their protection is unknown. We mapped the persistence of giant kelp forests across ten degrees of latitude in the Northeast Pacific Ocean and found that 7.7% of giant kelp is fully protected, with decreasing percentages from north to south. Sustainability goals should prioritize kelp mapping and monitoring, while protection and climate adaption targets should account for habitat dynamics.

## Main

Protected areas are a cornerstone for sustainability and biodiversity conservation^1^. As a result, the past decade has seen an increase in the area of marine and terrestrial ecosystems protected^2^, stimulated by international agreements that promote area-based conservation. The Convention on Biological Diversity (CBD) Aichi Target 11 and the Sustainable Development Goal 14^3,4^ aim to effectively protect at least 10% of ecologically representative coastal and marine areas by 2020, with increased aspirations to preserve 30% of oceans by 2030^5,6^. A central component of Aichi Target 11 is that protection includes a representative sample of coastal and marine habitats: many studies and national reports assess the representation of species and habitats such as corals, seagrass and mangroves^2,7,8^. However, some essential habitats like kelp forests remain neglected and information on their status and spatial distribution is largely lacking.

Kelp forests are one of the most productive^9^ ecosystems globally, comparable to coral reefs and terrestrial rainforests. Distributed along 25% of the world’s coastlines, they create a complex three-dimensional habitat, which sustains a diverse community of species^9,10^. However, extreme climatic events, overfishing, pollution, and other anthropogenic impacts threaten the capacity of these ecosystems to continue to produce goods and services worth billions of dollars to humanity^11,12^.

As marine heatwaves, hypoxic events, and other extreme episodes are becoming more frequent and severe^13^, ensuring the long-term persistence of species and ecosystems requires area-based conservation and adaptive strategies to address ongoing changes in climate and ocean chemistry^14^. One such strategy is protecting potential climate-refugia^15^, areas where the impacts of climate change may be less severe^16^. For dynamic ecosystems like kelp, that are highly variable on seasonal, annual, and decadal timescales^9^, it is critical to use long-term, large-scale datasets^17^ to understand their persistence, resilience and resistance^18^, and therefore identify potential climate refugia areas. If we map kelp forests and know patterns of persistence, we can prioritize their protection.

California, USA, and the Baja California Peninsula, Mexico, share the largest canopy-forming kelp forest species, the giant kelp *Macrocystis pyrifera*^19^ (henceforth “kelp”). This transboundary region has recently been subject to extreme marine heatwaves that decimated entire kelp forests^20,21^, threatening the outcomes of conservation efforts that established a network of marine protected areas in California^22^ and community-based marine reserves in Baja California^23^. Despite progress, recent reporting of marine habitat representation for both regions^7,8^ neglect kelp, and the conditions and location of kelp forests that are potential climate change refugia are unknown. How can countries meet post-2020 targets to adapt to climate change and protect 30% of marine habitats by 2030^5,6,14^, if no such information exists?

Here we map the distribution and persistence of highly dynamic *Macrocystis pyrifera* forests in the Northeast Pacific Ocean — spanning over ten degrees of latitude — using a 35 year satellite time series^24^. We quantified the representation of low, mid, and high persistent kelp found in two levels of protection (full and partial) across four distinct regions: Central and Southern California, and Northern and Central Baja California (see methods for details). Finally, we adjusted representation targets by calculating the additional area required to protect kelp that is expected to be present in any given year (see methods and supporting information for details).

Results show that, across the Northeast Pacific Ocean, 7.7% of kelp is fully protected and 3.9% is partially protected (Fig 1a). By level of persistence, 11.7% of highly persistent kelp is fully protected, with lower values for mid and low persistence (Fig 1a). By distribution, Central California has the highest amount of persistent kelp forest found in the Northeast Pacific Ocean (34.8%), while Northern Baja has the lowest (13.5%) (Fig 1b). In terms of protection by region, we found a decrease from north to south in the area coverage of fully protected kelp (Fig 1c, 2), being highest in Central (20.9%) and Southern California (8.4%) and lowest in Northern and Central Baja California (~1%) (Fig 2). We found a similar pattern for partially protected kelp (Fig 1c, 2). Central California also holds the highest percentage of highly protected persistent kelp (Fig 1c, 2).

**Fig. 1:**
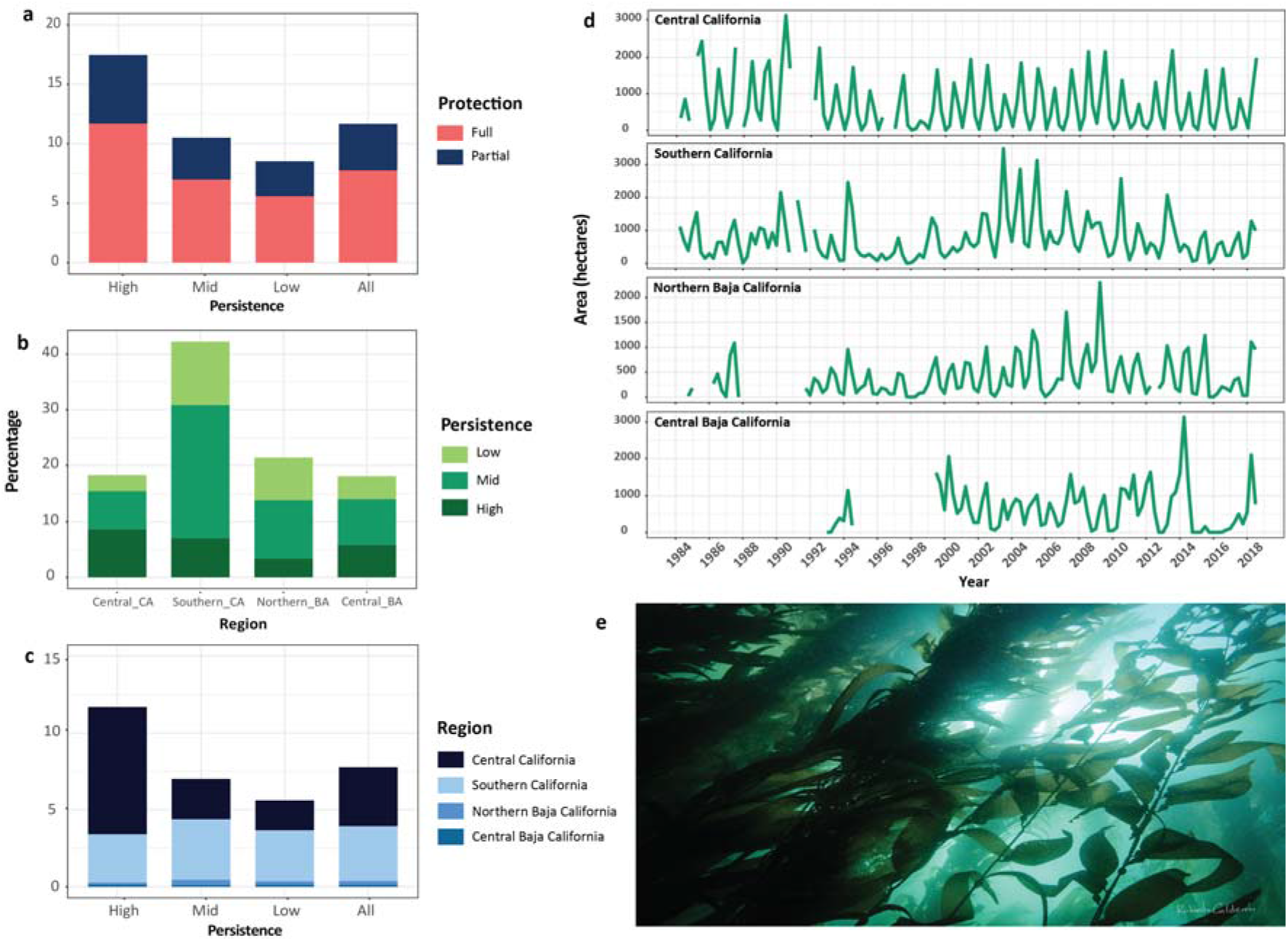
Protected kelp by level of persistence in the Northeast Pacific Ocean. Bar plots (left) show the percentage of the total kelp **a**, protected, **b**, distributed in each region, **c**, fully protected in each region. Time series (right) show **d**, the total area of kelp canopy for each region over the past 35 years; **e**, example of giant kelp ecosystem.

**Fig. 2:**
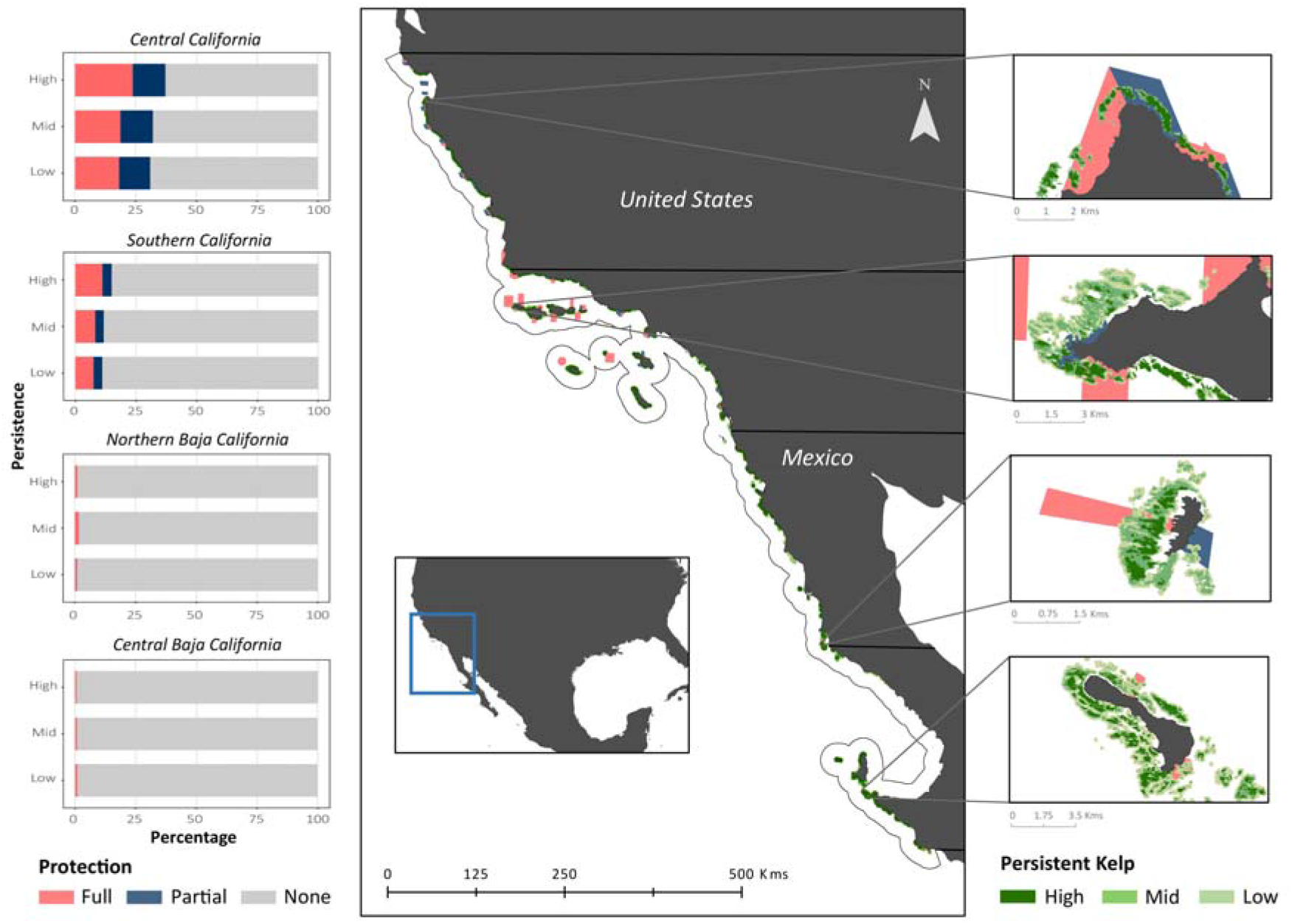
The spatial distribution of kelp by level of persistence and marine protected areas by level of protection. Left bar plots represent the percentage of persistent kelp protected per region. We provide fine-scale examples for each region.

We found an average persistence value of 0.43, which means that 43% of kelp distribution has kelp present in any year on average (see methods and Tab S1) in the Northeast Pacific Ocean, indicating that only 3.3% (instead of 7.7%) of kelp habitat expected to be present in any year is fully protected. The average persistence value ranged from 0.57 (Central California) to 0.37 (Northern Baja California), suggesting that fully protecting 10% of present kelp in each region requires, on average, an increase in the amount of kelp protected by over two-fold (see supporting information, Tab S1-2). However, these targets are smaller if we focus protection on highly persistent kelp, decreasing from 23.1 to 17.6% (Tab S1-2).

Marine reserves, or fully protected areas, are more effective than partial protection at conserving biodiversity^25^ and enhancing the resilience and adaptive capacity of ecosystems to climate impacts^26^. By fully protecting 7.7% of kelp, the Northeast Pacific Ocean appears to be approaching the CBD Aichi target 11 of effectively protecting 10% of coastal areas by 2020. However, Central California is the exception and additional investments are needed in the other regions. This is particularly urgent for Mexico, where 22% of its exclusive economic zone is protected^7^, but the extent of kelp protection in marine reserves in the coastal region of Baja California is extremely limited (~1%).

The uneven representation of persistent kelp in Baja California is of concern because the warm-distribution limit of the *Macrocystis pyrifera* is found here. This region is subject to episodes of higher sea surface temperatures and lower availability of nutrients, limiting kelp biomass and area^27^. Kelp found near their warm distribution limit are more impacted by extreme climatic events^21,27^, suggesting that future climate-driven impacts could significantly diminish the coverage of kelp in Baja California. Protection of persistent kelp in the region can minimize other local stressors, such as indirect negative effects of fishing (through removal of predators and release of herbivores that can over-graze kelp^28^), and build the resilience required for these ecosystems to adapt and persist in the face of future changes. For this reason, fully protecting the highly persistent kelp forests that are exhibiting high resilience to climate variability and extremes in Baja California is an urgent priority.

Unless the trend of increase in CO_2_ emissions is reversed, extreme climatic events are expected to become more frequent and severe in the following decades^29^, which will require science-based adaptation strategies in the Northeast Pacific Ocean. Protecting persistent kelp is one such strategy, but other measures will also be necessary, such as the restoration of degraded kelp, the identification of genetically resilient kelp stocks, and the management of other anthropogenic impacts not mitigated by marine reserves^11^. Importantly, we will need to test if persistent kelp acts as climate-refugia and understand the drivers and synergies (e.g., oceanographic features, human activities) which cause the high variability in local persistence (Fig 2), and how to integrate this information into the design of marine reserves.

Compared to less variable habitats like corals and mangroves, the highly dynamic nature of kelp forests^9^ (Fig 1d) poses unique challenges, rarely considered in conservation. Maps of kelp dynamics and persistence allow setting realistic and cost-effective habitat representation targets to protect kelp that is present in any given year. Not including this type of adjustments, can limit the amount of protected kelp that can provide the habitat structure for other community members.

Here, we illustrate how to map and identify potential climate-refugia for kelp and other highly dynamic habitats. We advise increased protection of highly persistent kelp given their potential climate-refugia attributes, wide-ranging ecosystem services and as a cost-effective approach to meet realistic area-based targets. Our effort should be scaled-up to map the global distribution and dynamics of kelp forests, which will require a globally coordinated effort. Only then, can countries assess their progress at meeting representation targets and support conservation and restoration actions for one of the world’s most productive ecosystems.

## Methods

The study area for this analysis encompasses the region where *Macrocystis pyrifera* is the dominant canopy kelp species in the Northeast Pacific Ocean. The region extends from Año Nuevo Island in the north (latitude ~37.1°), California, USA, to Punta Prieta in the south (latitude ~27°), Baja California Sur, Mexico. We mapped the distribution of giant kelp canopy and characterized persistence using a 30-m resolution satellite-based time series covering our entire study area (for a description of methods see^30^ and^21^ for the dataset). These data provide quarterly estimates of kelp canopy area across the study region from 1984-2018. We characterized kelp persistence as the average number of years occupied by kelp canopy (at least during one quarter in a year) in each pixel (n = 408906) for the past 35 years. Then, we used kelp persistence data (values of zero had no kelp, one occupied all years) to group pixels into three persistence classes. We classified pixels as low persistence in the 25^th^ percentile, with kelp found in less than 0.24 years. Mid persistence among the 25^th^ and 75^th^ percentile, with kelp found between 0.24 and 0.59 years. High persistence over the 75^th^ percentile, with kelp found over 0.59 years. To obtain the vectorial maps of kelp forest distribution for the three persistence levels, we rasterized the data points and converted them to polygons in ESRI ArcGIS Pro v10.8.

We obtained data on marine protected area location, boundary, and type for California from the National Oceanic and Atmospheric Administration (NOAA, 2020 version) and for community-based marine reserves in the Baja California Peninsula from Comunidad y Biodiversidad, an NGO that has been supporting the local fishing cooperatives in establishing the voluntary reserves. We performed a spatial overlay analysis to estimate the representation of kelp habitats in marine protected areas. We performed the analysis using ESRI ArcGIS Pro v10.8, calculating coverage through spatial intersections of two marine protected area categories (no-take and multiple-use) and kelp forest persistence (high, mid, and low) for our region. We combined and merged marine protected areas based on the two levels of protection: no-take areas are the most restrictive type where all extractive uses are prohibited (full protection), and multiple-use areas where some restrictions apply to recreational and commercial fishing (partial protection). We divided our region in four areas, Central and Southern California, and Northern and Central Baja California. These four regions represent distinct biogeographic areas^31^ were species composition vary because of oceanographic forcing, or geographic borders (USA and Mexico border). We conducted the analysis for the entire region and separately for each of the four regions.

Finally, we estimate a multiplier required to ensure we are meeting representation targets for protecting kelp forest habitat that is present, rather than just its potential distribution (see supporting information). We define present kelp as the probability that a pixel will have kelp in any given year, thus maintaining the habitat structure they provide, and potential kelp distribution as any pixel covered by kelp at least once in the time series.

## Supporting information

Methods to estimate the representation target for present kelp

## Acknowledgements

N.A.-D acknowledges support from the Fundación Bancaria ‘la Caixa’ under the Postgraduate Fellowship (LCF/BQ/AA16/11580053), from the University of Queensland under the Research Training Scholarship, and from the Estate Winifred Violet Scott for a research grant. F.M. acknowledges the support of the US NSF (OCE 1736830).

## Contributions

N.A.-D conceived the idea with inputs from K.C.C, H.P.P, and F.M. N.A.-D conducted the spatial analysis, K.C.C led the kelp mapping with the support of K.C and T.B. N.A.-D wrote the manuscript with editorial input from all other authors.

## Corresponding author

Correspondence to Nur Arafeh Dalmau

## Ethics declaration

The authors declare no competing interests

## Notes

### Competing Interest Statement

The authors have declared no competing interest.

## References

1 Lester, S. E. et al. Biological effects within no-take marine reserves: a global synthesis. Marine Ecology Progress Series 384, 33–46 (2009).

2 Maxwell, S. L. et al. Area-based conservation in the twenty-first century. Nature 586, 217–227 (2020).

3 Diversity., S. o. t. C. o. B. COP 10 Decision X/2: strategic plan for biodiversity 2011–2020 (2010).

4 Ga, U. Transforming our world: the 2030 Agenda for Sustainable Development. Division for Sustainable Development Goals: New York, NY, USA (2015).

5 IUCN. Motion 053: increasing marine protected area coverage for effective marine biodiversity conservation. (2016).

6 Diversity, C. O. B. (2020).

7 CONANP. Resiliencia Áreas Naturales Protegidas Soluciones naturales a retos globales. (2019).

8 NOAA. Marine Protected Areas 2020: Building Efective Conservation Networks (2020).

9 Schiel, D. R. & Foster, M. S. The biology and ecology of giant kelp forests. (Univ of California Press, 2015).

10 Wernberg, T., Krumhansl, K., Filbee-Dexter, K. & Pedersen, M. F. in World seas: an environmental evaluation 57–78 (Elsevier, 2019).

11 Arafeh-Dalmau, N. et al. Marine heat waves threaten kelp forests. Science 367, 635–635 (2020).

12 Smale, D. A.. et al. Marine heatwaves threaten global biodiversity and the provision of ecosystem services. Nature Climate Change 9, 306–312 (2019).

13 Frölicher, T. L., Fischer, E. M. & Gruber, N. Marine heatwaves under global warming. Nature 560, 360–364 (2018).

14 Roberts, C. M., O’Leary, B. C. & Hawkins, J. P. Climate change mitigation and nature conservation both require higher protected area targets. Philosophical Transactions of the Royal Society B 375, 20190121 (2020).

15 Wilson, K. L., Tittensor, D. P., Worm, B. & Lotze, H. K. Incorporating climate change adaptation into marine protected area planning. Global Change Biology (2020).

16 Keppel, G. et al. The capacity of refugia for conservation planning under climate change. Frontiers in Ecology and the Environment 13, 106–112 (2015).

17 Hughes, B. B. et al. Long-term studies contribute disproportionately to ecology and policy. BioScience 67, 271–281 (2017).

18 O’Leary, J. K. et al. The resilience of marine ecosystems to climatic disturbances. BioScience 67, 208–220 (2017).

19 Aburto-Oropeza, O. et al. Harnessing cross-border resources to confront climate change. Environmental Science & Policy 87, 128–132 (2018).

20 Arafeh-Dalmau, N. et al. Extreme marine heatwaves alter kelp forest community near its equatorward distribution limit. Frontiers in Marine Science 6, 499 (2019).

21 Cavanaugh, K. C., Reed, D. C., Bell, T. W., Castorani, M. C. & Beas-Luna, R. Spatial variability in the resistance and resilience of giant kelp in southern and Baja California to a multiyear heatwave. Frontiers in Marine Science 6, 413 (2019).

22 Kirlin, J. et al. California’s Marine Life Protection Act Initiative: supporting implementation of legislation establishing a statewide network of marine protected areas. Ocean & Coastal Management 74, 3–13 (2013).

23 A.C., C. y. B. Reservas marinas totalmente protegidas en México (2005-2016). Comunidad y Biodiversidad, A.C., Guaymas, Sonora, México. (2018).

24 Bell, T. W., Cavanaugh, K. C. & Siegel, D. A. SBC LTER: time series of quarterly NetCDF files of kelp biomass in the canopy from Landsat 5, 7 and 8, 1984-2016 (ongoing). Environmental Data Initiative (2020).

25 Sala, E. & Giakoumi, S. No-take marine reserves are the most effective protected areas in the ocean. ICES Journal of Marine Science 75, 1166–1168 (2018).

26 Roberts, C. M. et al. Marine reserves can mitigate and promote adaptation to climate change. 114, 6167–6175 (2017).

27 Hernandez-Carmona, G., Robledo, D. & Serviere-Zaragoza, E. Effect of nutrient availability on Macrocystis pyrifera recruitment and survival near its southern limit off Baja California. Botanica Marina 44, 221–229 (2001).

28 Ling, S., Johnson, C., Frusher, S. & Ridgway, K. Overfishing reduces resilience of kelp beds to climate-driven catastrophic phase shift. Proceedings of the National Academy of Sciences 106, 22341–22345 (2009).

29 Oliver, E. C. et al. Projected marine heatwaves in the 21st century and the potential for ecological impact. Frontiers in Marine Science 6, 734 (2019).

30 Bell, T. W., Allen, J. G., Cavanaugh, K. C. & Siegel, D. A. Three decades of variability in California’s giant kelp forests from the Landsat satellites. Remote Sensing of Environment 238, 110811 (2020).

31 Blanchette, C. A. et al. Biogeographical patterns of rocky intertidal communities along the Pacific coast of North America. Journal of Biogeography 35, 1593–1607 (2008).

