## Supplementary material for "Uneven protection of persistent giant kelp forest in the Northeast Pacific Ocean": Methods to estimate the representation target for present kelp

### Supplementary information

### Methods to estimate the representation target for present kelp

#### 1) Fixed targets

In this method, we estimate a multiplier required to meet representation targets for present kelp rather than potential kelp distribution. We estimated persistence values for each pixel (n = 408906), as the percentage of years (at least during one quarter in a year) occupied by kelp forest in that pixel for the past 35 years (see method section for detail). In other words, the probability of kelp forest occurrence in that pixel in any given year. We define a pixel to be potential kelp habitat if the satellite identified kelp forest in that pixel in at least one quarter of a year in the time series. A pixel with zero value means the satellite never found kelp forest and thus it is not potential kelp; while a value of one means, it detected kelp forest for all years. We define the probability of persistence as the average years that kelp forest occupied any given pixel with potential kelp habitat in the time series as *P*:

$$P = \frac{\sum_{i=1}^{n} O_i}{n},$$

where $O_i$ is the fraction of years occupied by potential kelp habitat for pixel *i* and *n* the number of potential kelp habitat pixels. The probability of persistence (*P*) is equivalent to the probability of any given pixel to have present kelp in any year in a given region. We can now estimate the representation of present kelp ($R_p$):

$$R_p = TP,$$

where *T* is the representation target and *P* the probability of persistence. $R_p$ gives as an estimate of the percentage of present kelp habitat protected. We can now apply a multiplier, *M*:

$$M = \frac{1}{P},$$

which adjusts the representation target ($T_a$):

$$T_a = \text{TM} * 100,$$

ensuring the representation of present kelp ($R_p$) to be equivalent to the representation target (*T*).

2. **Unfixed targets**

The previous approach uses fixed representation targets without accounting for the classification of kelp based on their persistence (see methods in manuscript). However, we can adjust representation targets for specific persistence classes. As an example, we can only adjust representation target for highly persistence kelp. We can then use the previous equation for each level of persistence (low, mid, high), leaving constant the representation target (*T*) for low and mid persistence, and estimate the adjusted representation target for highly persistence kelp ($T_h$):

$$Tn = R_l n_l + R_m n_m + R_h n_h ,$$

$$Tn = TP_l n_l + TP_m n_m + T_h P_h n_h ,$$

$$T_h = \frac{T\,(n - P_l n_l + P_m n_m)}{P_h n_h},$$

where $R_l$, is the representation of low, $R_m$ mid, and $R_h$ high present kelp. Then $P_l$ is the probability of low, $P_m$ for mid, and $P_h$ for high persistence kelp. Finally, *n* is the number of potential kelp habitat pixels, $n_l$ is the number of pixels with low, $n_m$ with mid, and $n_h$ with high persistence kelp. We can then estimate the multiplier required to adjust representation targets of highly persistence kelp $M_h$:

$$M_h = \frac{T_h}{T}$$

**<u>Example</u>**

1) **Fixed targets**

We provide an example by estimating the representation of present kelp and the multiplier (*M*) required to represent 10% of present kelp ($R_p$) in the Northeast Pacific Ocean:

$$R_p = 0.1 * 0.43 * 100 ,$$

where the probability of persistence (*P*) is 0.43 and the representation target (*T*) is 0.1. By protecting 10% of potential kelp habitat, only 4.3% of present kelp is protected in the Northeast Pacific Ocean. We can now we estimate the multiplier (*M*):

$$M = \frac{1}{0.43},$$

which suggest that we need to apply a multiplier (M) of 2.31 to protect 10% of present kelp in the Northeast Pacific Ocean. Finally, we can adjust the representation target ($T_a$):

$$T_a = 0.1 * 2.31 * 100\,,$$

which suggests that we need to protect 23.1% of potential kelp habitat if we account for persistence.

**2) Unfixed targets**

We also provide an example by estimating the adjusted representation target of highly persistence kelp ($T_h$) required to represent 10% of present kelp in the Northeast Pacific Ocean:

$$T_h = \frac{0.1(408906 - 0.16 * 99477 - 0.42 * 207744)}{0.74 * 101685} * 100\,,$$

which suggests that we need to protect 40.7% of highly persistence kelp to meet representation target (T) and apply a multiplier for highly persistence kelp ($M_h$):

$$M_h = \frac{0.407}{0.1}\,,$$

of 4.07. See values from Table S1 (next).

**TABLES**

**Table S1-** Probability of persistence (*P*) and number of potential kelp pixels (*n*) for each region and for the Northeast Pacific Ocean combined without classifying based on their persistence. Low capital letters for the probability of persistence (*P*) and the number of potential kelp pixels (*n*) represent values based on low ($_l$), mid (m), and high ($_h$) persistence.

| **Region** | $P$ | $n$ | $P_l$ | $n$ | $P_m$ | $n$ | $P_h$ | $n$ |
|---|---|---|---|---|---|---|---|---|
| Central California | 0.57 | 69,633 | 0.17 | 8,840 | 0.41 | 27,555 | 0.81 | 33,238 |
| Southern California | 0.40 | 177,745 | 0.16 | 43,831 | 0.41 | 102,991 | 0.71 | 30,923 |
| Northern Baja California | 0.37 | 87,645 | 0.14 | 30,396 | 0.43 | 43,479 | 0.70 | 13,770 |
| Central Baja California | 0.46 | 73,883 | 0.16 | 16,410 | 0.43 | 33,719 | 0.71 | 23,754 |
| Northeast Pacific Ocean | 0.43 | 408,906 | 0.16 | 99,477 | 0.42 | 207,744 | 0.74 | 101,685 |

**Table S2-** Adjusted representation targets for fixed (*M*) and unfixed ($M_h$) present kelp and area requirements for each region and for the Northeast Pacific Ocean combined.

| **Region** | M | Area | *$M_h$ | Area |
|---|---|---|---|---|
| Central California | 1.75 | 17.5% | 2.08 | 15.4% |
| Southern California | 2.5 | 25% | 5.60 | 18.5% |
| Northern Baja California | 2.71 | 27.1% | 6.34 | 19.1% |
| Central Baja California | 2.17 | 21.7 | 3.38 | 17.7% |
| Northeast Pacific Ocean | 2.31 | 23.1% | 4.07 | 17.6% |

*we kept constant the representation target for low and mid persistence (10%), and estimated the multiplier for highly persistent kelp needed to meet the representation target of 10%.
